# The Omicron variant is highly resistant against antibody-mediated neutralization – implications for control of the COVID-19 pandemic

**DOI:** 10.1101/2021.12.12.472286

**Authors:** Markus Hoffmann, Nadine Krüger, Sebastian Schulz, Anne Cossmann, Cheila Rocha, Amy Kempf, Inga Nehlmeier, Luise Graichen, Anna-Sophie Moldenhauer, Martin S. Winkler, Martin Lier, Alexandra Dopfer-Jablonka, Hans-Martin Jäck, Georg M. N. Behrens, Stefan Pöhlmann

## Abstract

The rapid spread of the SARS-CoV-2 Omicron variant suggests that the virus might become globally dominant. Further, the high number of mutations in the viral spike-protein raised concerns that the virus might evade antibodies induced by infection or vaccination. Here, we report that the Omicron spike was resistant against most therapeutic antibodies but remained susceptible to inhibition by Sotrovimab. Similarly, the Omicron spike evaded neutralization by antibodies from convalescent or BNT162b2-vaccinated individuals with 10- to 44-fold higher efficiency than the spike of the Delta variant. Neutralization of the Omicron spike by antibodies induced upon heterologous ChAdOx1/BNT162b2-vaccination or vaccination with three doses of BNT162b2 was more efficient, but the Omicron spike still evaded neutralization more efficiently than the Delta spike. These findings indicate that most therapeutic antibodies will be ineffective against the Omicron variant and that double immunization with BNT162b2 might not adequately protect against severe disease induced by this variant.

## INTRODUCTION

Vaccination is considered key to ending the devastating COVID-19 pandemic. However, inequities in vaccine distribution and the emergence of new SARS-CoV-2 variants threaten this approach. Several SARS-CoV-2 variants of concern (VOC) have emerged in the recent year and the Delta variant (B.1.617.2) is currently dominating the pandemic (Harvey et al., 2021b; Tao et al., 2021). These VOC exhibit increased transmissibility and/or immune evasion, traits that have been linked to mutations in the viral spike protein (S) (Harvey *et al*., 2021b; Tao *et al*., 2021).

The coronavirus S protein facilitates viral entry into host cells and constitutes the central target for antibodies that neutralize the virus. Mutations in the N-terminal domain (NTD), which contains an antigenic supersite (McCallum et al., 2021), and the receptor binding domain (RBD), which binds to the ACE2 receptor (Hoffmann et al., 2020b; Zhou et al., 2020), can confer neutralization resistance by altering epitopes of neutralizing antibodies. In contrast, it is less well understood which mutations in spike increase transmissibility and which mechanisms are responsible, although it is well established that mutation D614G increases viral transmission and promotes ACE2 engagement (Hou et al., 2020; Korber et al., 2020; Mansbach et al., 2021; Plante et al., 2021; Zhou et al., 2021).

A novel VOC, the Omicron variant (Pango lineage B.1.1.529 and sublineages BA.1 and BA.2), was recently identified in South Africa and its emergence was associated with a steep increase in cases and hospitalizations (Abdullah, 2021). The Omicron variant was imported into several European, African and Asian countries as well as the USA via infected air travelers (Abbasi, 2021; Graham, 2021; Gu et al., 2021; Petersen et al., 2021). In the United Kingdom, local transmission events were reported (Company, 2021) with case numbers doubling every two to three days (Torjesen, 2021). The S protein of the Omicron variant harbors an unusually high number of mutations, which might increase immune evasion and/or transmissibility. Indeed, a recent study suggested that the Omicron variant is more adept at infecting convalescent individuals as compared to previously circulating variants (Abdullah, 2021; Pulliam et al., 2021). Thus, the Omicron variant constitutes a rapidly emerging threat to public health and might undermine global efforts to control the COVID-19 pandemic. However, the susceptibility of the Omicron variant to antibody-mediated neutralization remains to be analyzed.

Here, we report that the Omicron S protein evades antibodies with up to 44-fold higher efficiency than the spike of the Delta variant, rendering therapeutic antibodies ineffective and likely compromising protection by antibodies induced upon infection or vaccination with two doses of BNT162b2 (BNT).

## RESULTS

### The NTD and RBD of the Omicron spike are highly mutated

The first sequences of the Omicron variant (BA. 1) were deposited into the GISAID (Global Initiative on Sharing All Influenza Data) database between 22 and 23 of November 2021 (Figure 1A). These sequences were obtained from patients in Botswana and South Africa as well as from a traveler returning back to Hong Kong from South Africa. Subsequently, the number of deposited Omicron sequences rapidly increased (Figure 1A) and the virus was also detected in Europe, Asia and the USA due to infected air travelers (Figure 1B). Analysis of the Omicron genomic sequence revealed striking differences as compared to other known SARS-CoV-2 variants, suggesting extensive, independent evolution, potentially in an isolated human population, immunocompromised patients or an unknown animal species (Kupferschmidt, 2021).

**Figure 1:**
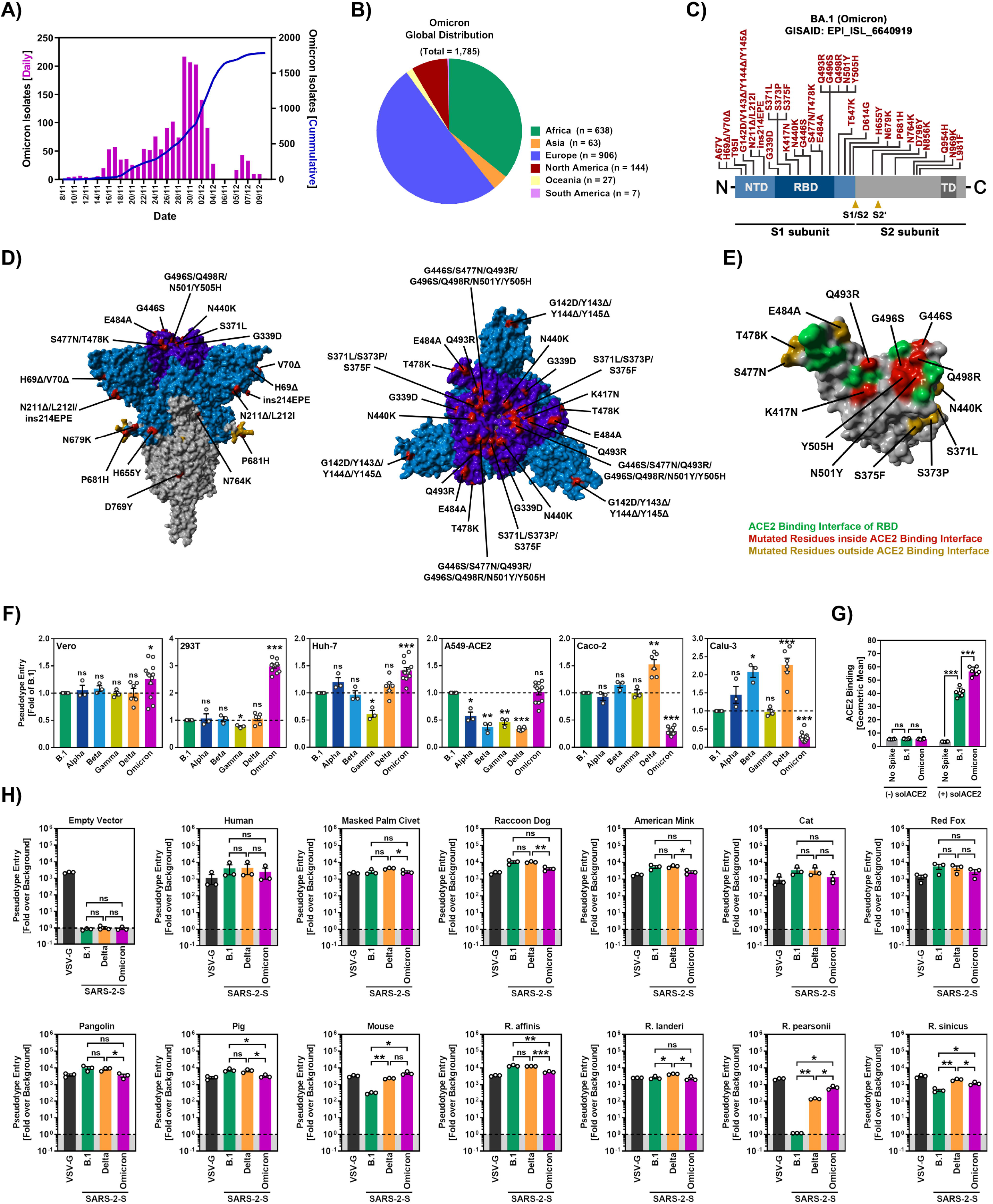
The Omicron spike displays cell line-specific differences in cell tropism, bind ACE2 with increased efficiency and utilizes a broad range of animal ACE2 orthologues as receptor. (A) Epidemiology of SARS-CoV-2 Omicron (BA. 1) variant. Purple bars indicate the number of newly reported isolates per day while the blue line show the cumulative number of isolates (as of 11.10.2021; based on data deposited in the GISAID database). (B) Global distribution of SARS-CoV-2 Omicron variant. Sequences retrieved from the GISAID database were grouped based on the continent where they were detected (as of 11.10.2021). (C) Schematic overview and domain organization of the S proteins of SARS-CoV-2 Omicron variant. Mutations compared to the SARS-CoV-2 Wuhan-Hu-1 isolate are highlighted in red. Abbreviations: NTD, N-terminal domain; RBD, receptor-binding domain; TD, transmembrane domain. (D) Location of Omicron-specific mutations in the context of the trimeric spike protein. Color code: light blue, S1 subunit with RBD in dark blue; gray, S2 subunit; orange, S1/S2 and S2’ cleavage sites; red, mutated amino acid residues (compared to the S protein of the SARS-CoV-2 Wuhan-Hu-1 isolate) (E) Location of Omicron-specific mutations in the context of the RBD. Color code: gray, RBD; green, RBD residues that interact ACE2; red, ACE2-interacting RBD residues that are mutated in Omicron spike: orange, non-ACE2-interacting RBD residues that are mutated in Omicron spike (compared to the S protein of the SARS-CoV-2 Wuhan-Hu-1 isolate) (F) Particles bearing the indicated S proteins were inoculated on different cell lines and S protein-driven cell entry was analyzed at 16-18 h postinoculation by measuring the activity of virus-encoded firefly luciferase in cell lysates. Presented are the average (mean) data from three to twelve biological replicates (each conducted with four technical replicates) in which S protein-driven cell entry was normalized against B.1 (set as 1). Error bars indicate the standard error of the mean (SEM). (G) Binding of soluble ACE2 to B.1 or Omicron spike proteins. 293T-cells expressing the indicated S protein (or no S protein) following transfection, were sequentially incubated with soluble ACE2 harboring a C-terminal Fc-tag derived from human IgG and AlexaFlour-488-conjugated anti-human antibody. Finally, ACE2 binding was analyzed by flow cytometry. Cells incubated only with secondary antibody served as controls. Presented is the average (mean) geometric mean channel fluorescence from six biological replicates (each conducted with single samples). Error bars indicate the standard deviation. (H) Particles bearing the indicated S proteins or VSV-G were inoculated on BHK-21 cells that were previously transfected to express the indicated ACE2 orthologues or empty vector. S protein-driven cell entry was analyzed as described in panel F. Presented are the average (mean) data from three biological replicates (each conducted with four technical replicates) in which signals obtained from particles bearing no viral glycoprotein (indicated by dashed line) were used for normalization (set as 1). Error bars indicate the standard error of the mean (SEM). Panels F-H: Statistical significance of differences between individual groups was assessed by two-tailed Students t-test (p > 0.05, not significant [ns]; p < 0.05, *; p < 0.01, **; p < 0.001, ***).

The Omicron spike exhibits 37 mutations as compared to the Wuhan-Hu-1 spike (Figure 1C-D). 13 of these changes are unique while the remaining changes are known from variants of interest or concern (Haseltine, 2021). Specifically, 11 mutations, including 6 deletions and 1 insertion, are located in the NTD and mutations N211Δ and ins214EPE are unique. Some of the deletions are found in other VOC and might reduce antibody binding or increase spike expression (Harvey *et al*., 2021b; Li et al., 2021b; Liu et al., 2021; Meng et al., 2021; Wang et al., 2021; Wheatley et al., 2021). Another 15 mutations are found in the RBD (Figure 1C-E), of which G339D, S371L, S373P and S375F are unique, and several were previously shown to modulate ACE2 binding and/or antibody evasion (Harvey *et al*., 2021b; Li *et al*., 2021b; Liu *et al*., 2021; Wang *et al*., 2021; Wheatley *et al*., 2021). Further, 5 mutations reside between RBD and S1/S2 site, including the unique mutation T547K and mutation P681H that may modulate cleavage at the S1/S2 site by the host protease furin (Hoffmann et al., 2020a; Zhang et al., 2021a) (Figure 1C and D). Finally, five mutations are present in the S2 subunit, all of them unique (Haseltine, 2021).

### Omicron spike binds human ACE2 and mediates efficient entry into cell lines

For analysis of host cell entry, we employed vesicular stomatitis virus (VSV) particles pseudotyped with SARS-CoV-2 S proteins. These pseudotyped particles adequately mimic key features of SARS-CoV-2 entry into target cells, including receptor and protease choice and neutralization by antibodies (Hoffmann *et al*., 2020b; Riepler et al., 2020; Schmidt et al., 2020).

We first asked whether the Omicron spike differs from the spike of other VOCs regarding target cell choice and entry efficiency. The spike from SARS-CoV-2 B.1 (which is identical to the S protein of the Wuhan-Hu-1 isolate except for the presence of mutation D614G) was analyzed in parallel, since this virus circulated early in the pandemic and does not harbor mutations found in the S proteins of VOCs. For the analysis of cell tropism, we employed the cell lines Vero (African green monkey, kidney), 293T (human, kidney), A549 (human, lung) engineered to express ACE2, Huh-7 (human, liver), Caco-2 (human, colon) and Calu-3 (human, lung) cells. All cell lines were highly susceptible to entry driven by VSV-G and SARS-CoV-2 B.1 spike (Figure 1F and supplemental figure S1). Further, all VOC S proteins facilitated robust entry into the cell lines analyzed but subtle differences were noted. Thus, the Delta spike mediated increased entry into Calu-3 and Caco-2 cells, in keeping with published data (Arora et al., 2021b), while the Omicron spike facilitated increased entry into Vero, Huh-7 and particularly 293T cells (Figure 1F). Further, B.1 and Omicron spike mediated comparable entry into A549-ACE2 cells while entry driven by the other S protein was less efficient (Figure 1F). Finally, the Omicron spike bound efficiently to human ACE2 (Figure 1G) and used ACE2 for host cell entry (see below), indicating that the mutations in the RBD do not compromise ACE2 interactions. In sum, the Omicron spike binds to ACE2 and mediates robust entry into diverse ACE2-positive cell lines.

### The Omicron spike can efficiently use ACE2 orthologues from different animal species for cell entry

We next asked whether the Omicron spike can use human ACE2 and ACE2 orthologues from different animal species, including horseshoe bats, masked palm civet, raccoon dog and pangolin, for entry into target cells. Entry driven by the B. 1 and Delta spikes as well as VSV-G served as controls. The expression of the ACE2 orthologues did not modulate entry driven by VSV-G but in most cases allowed robust and roughly comparable entry driven by the B.1, Delta and Omicron spikes (Figure 1H). Two exceptions were noted. Murine ACE2 was used more efficiently by the Delta spike as compared to the B.1 spike and supported entry driven by the Omicron spike with the highest efficiency (Figure 1H). Finally, B.1 spike was unable to use ACE2 from Pearson’s horseshoe bat (*Rhinolophus pearsonii*) for entry while Delta and particularly Omicron spike used this receptor with high efficiency (Figure 1H). Collectively, these data reveal broad usage of ACE2 orthologues by Omicron spike for host cell entry, which might hint towards high zoonotic potential.

### Omicron spike is inhibited by soluble ACE2 but is resistant against most antibodies used for COVID-19 treatment

We next asked whether the Omicron spike can be inhibited by soluble ACE2, which binds to the RBD, blocks host cell entry and is currently being developed for COVID-19 therapy (Monteil et al., 2020). Soluble ACE2 did not modulate entry driven by VSV-G but robustly and comparably inhibited entry driven by B.1, Delta and Omicron spike (Figure 2A), indicating that soluble ACE2 should be suitable for treatment of patients infected with the Omicron variant.

**Figure 2:**
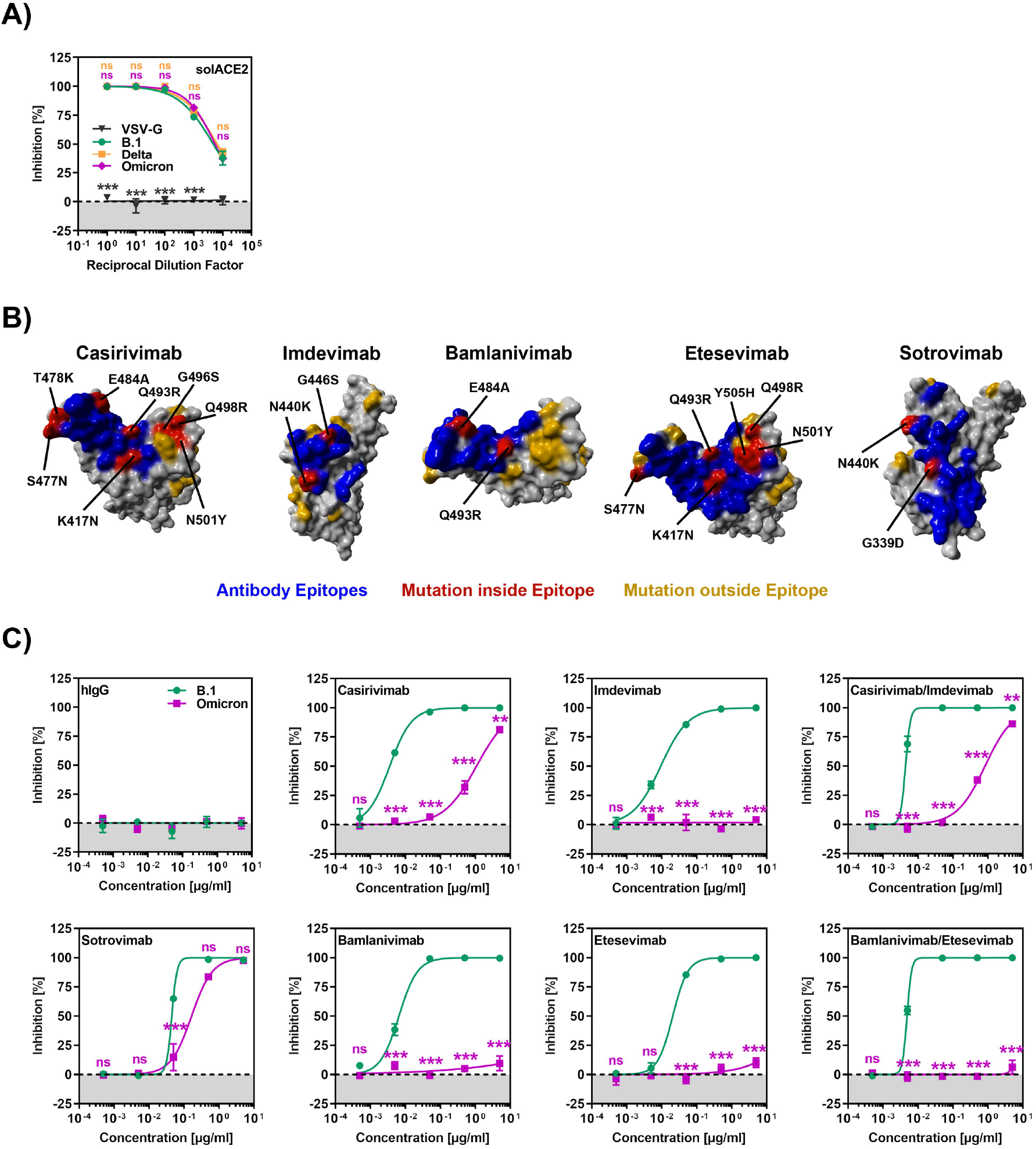
The Omicron spike evades neutralization by four out five monoclonal antibodies and is efficiently inhibited by soluble ACE2. (A) Inhibition of S protein-driven cell entry by soluble ACE2. Particles harboring the indicated S proteins were preincubated (30 min, 37 °C) with different dilutions of soluble ACE2 before being inoculated on Vero cells. S protein-driven cell entry was analyzed as described in Fig. 1F. Presented are the average (mean) data from three biological relocates (each conducted with four technical replicates). (B) Location of Omicron-specific mutations in the context of the RBD epitopes targeted by Casirivimab, Imdevimab, Bamlanivimab, Etesevimab and Sotrovimab. Color code: gray, RBD, blue epitope targeted by the antibody; red, Omicron-specific mutations within the epitope; orange, Omicron-specific mutations outside the epitope (C) Omicron spike is resistant against four out of five monoclonal antibodies used for treatment of COVID-19 patients. Particles harboring the indicated S proteins were preincubated (30 min, 37 °C) with different concentrations of recombinant monoclonal antibody before being inoculated on Vero cells. S protein-driven cell entry was analyzed as described in Fig. 1F. Presented are the average (mean) data from three biological relocates (each conducted with four technical replicates). Panels A and C: Statistical significance of differences between individual groups was assessed by two-way analysis of variance with Dunnet’s (panel A) or Sidak’s (panel C) post-hoc test (p > 0.05, not significant [ns]; p < 0.05, *; p < 0.01, **; p < 0.001, ***).

Several recombinant, neutralizing monoclonal antibodies were identified that inhibit SARS-CoV-2 infection and cocktails of Casirivimab and Imdevimab (REGN-COV2)(Weinreich et al., 2021) and Etesevimab and Bamlanivimab (Eli Lilly)(Weinreich *et al*., 2021) are currently used for COVID-19 therapy. In addition, the antibody Sotrovimab was shown to inhibit SARS-CoV-2 and related viruses and was found to protect patients from COVID-19 (Gupta et al., 2021). Since the Omicron spike harbors several mutations within the structures that are recognized by these antibodies (Figure 2B), we investigated whether the antibodies were still able to neutralize the Omicron spike. All antibodies inhibited entry driven by the B.1 spike in a robust and concentration dependent manner while a control immunoglobulin was inactive (Figure 2C). In contrast, entry driven by the Omicron spike was fully resistant against Bamlanivimab, Etesevimab and Imdevimab and largely resistant against Casirivimab. In agreement with these findings, a cocktail of Bamlanivimab and Etesevimab failed to inhibit entry mediated by Omicron spike while inhibition by a cocktail of Casirivimab and Imdevimab was inefficient (Figure 2C). In contrast, Sotrovimab was active against Omicron spike, although inhibition was slightly less efficient than that measured for B. 1 spike (Figure 2C). In sum, the Omicron spike is resistant against most antibodies used for COVID-19 treatment.

### The Omicron spike evades neutralization by antibodies induced upon infection and BNT vaccination with high efficiency

The resistance against several antibodies used for COVID-19 therapy suggested that the Omicron spike might also evade antibodies induced upon infection and vaccination. Indeed, sera/plasma collected within 2 months of convalescence from mild or severe COVID-19 inhibited entry driven by the Omicron spike 80–fold less efficiently as compared to the B. 1 spike and 44-fold less efficiently as compared to Delta spike, with 9 out of 17 sera tested being unable to neutralize particles bearing Omicron spike (Figure 3A and supplemental figure S2). The samples were collected in Germany during the first COVID-19 wave (supplemental table S1), when neither the Alpha nor the Delta variant predominated, suggesting the antibodies raised against the virus circulating at the beginning of the pandemic offer little to no protection against the Omicron variant.

**Figure 3:**
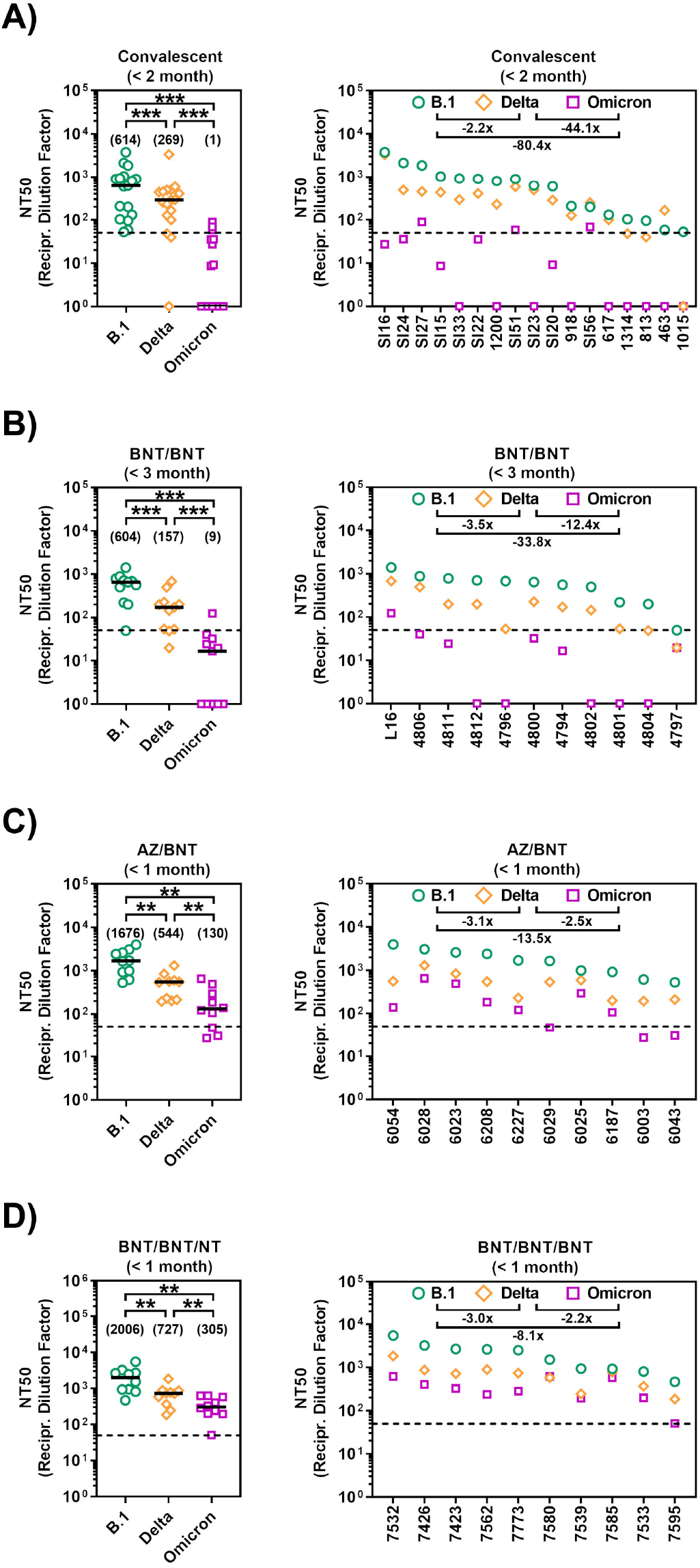
The Omicron spike shows high resistance against antibodies elicited upon infection or vaccination. (A) Particles bearing the indicated S proteins were preincubated (30 min, 37 °C) with different dilutions of convalescent sera/plasma (n = 17) before being inoculated on Vero cells. S protein-driven cell entry was analyzed as described in Fig. 1F. (B) The experiment was performed as described in panel A but sera from BNT/BNT-vaccinated individuals were analyzed at an early (< 3 month after receiving the second jab, n = 11) or late phase (> 6 month after receiving the second jab, n = 9) after vaccination. (C) The experiment was performed as described in panel A but sera from AZ/BNT-vaccinated individuals were analyzed (n = 10). (D) The experiment was performed as described in panel A but sera from BNT/BNT/BNT-vaccinated individuals were analyzed (n = 10). All panels: Shown are the reciprocal serum/plasma dilution factors that caused a 50 % reduction in S protein-driven cell entry (Neutralization titer 50, NT50). Left panels show individual NT50 values clustered per SARS-CoV-2 variant (Black lines and numerical values in brackets indicate median NT50 values) whereas right panels show serum/plasma-specific NT50 values ranked from highest to lowest (based on NT50 for B.1; numerical values indicate the median fold change in neutralization between individual SARS-CoV-2 variants). Dashed lines indicate the lowest serum dilution tested. Statistical significance of differences between individual groups was assessed by Wilcoxon matched-pairs signed rank test (p > 0.05, not significant [ns]; p < 0.05, *; p < 0.01, **; p < 0.001, ***).

The mRNA-based vaccine BNT efficiently protects against COVID-19 (Polack et al., 2020) and is frequently used in Europe and the USA. Sera collected within three months after the second dose of BNT (supplemental table S2) inhibited entry by the Omicron spike with 34-fold lower efficiency as compared to the B.1. spike and with 12-fold lower efficiency as compared to Delta spike (Figure 3B and supplemental figure S2). These results suggest that two immunizations with BNT, which can provide more than 90% protection from severe disease upon infection with the Delta variant (Chemaitelly et al., 2021), might be markedly less effective against the Omicron variant.

### Evidence that heterologous and booster vaccination may provide improved protection against the Omicron variant

We next explored whether strategies known to increase production of neutralizing antibodies relative to BNT/BNT immunization might afford better protection against the Omicron variant. Heterologous vaccinations with a first of dose of ChAdO1-nCoV-19/AZD1222 (AZ) (Voysey et al., 2021) and a second dose of BNT was shown to induce higher neutralizing antibody titers as compared to the corresponding homologous vaccinations (Barros-Martins et al., 2021; Behrens et al., 2021). Sera collected within 1 month after heterologous AZN/BNT vaccination (supplemental table S2) exhibited higher neutralizing activity as compared to sera collected within 3 months after BNT/BNT vaccination (Figure 3C). Inhibition of Omicron spike-driven entry by sera from AZ/BNT vaccinated individuals was 14-fold less efficient as compared to B.1. spike but only 3-fold less efficient relative to Delta spike (Figure 3C). Further, inhibition of Omicron spike by sera collected within 1 months of AZ/BNT vaccination was comparable to inhibition of Delta spike by sera obtained within 3 months after BNT/BNT vaccination (Figure 3B and C), a time interval during which the vaccine provides more than 90% protection from severe COVID-19.

A third immunization with BNT increases protection against infection by 10-fold as compared to two doses of BNT (Bar-On et al., 2021). Sera from BNT/BNT/BNT immunized donors collected within 1 month after the third dose (supplemental table S2) had slightly higher neutralizing titers as sera obtained within the same time interval from donors immunized with AZ/BNT (Figure 3C-D). Sera from BNT/BNT/BNT immunized individuals inhibited entry driven by Omicron spike with 8-fold reduced efficiency as compared to B.1 spike and 2-fold reduced efficiency as compared to Delta spike (Figure 3D). Further, inhibition of Omicron spike by sera collected within 1 months after BNT/BNT/BNT immunization was moderately more efficient as compared to inhibition of Delta spike by sera obtained within 3 months after BNT/BNT vaccination (Figure 3B and D). These results suggest that heterologous AZ/BNT as well as homologous BNT/BNT/BNT immunization might provide better protection against the Omicron variant as compared to BNT/BNT immunization.

## DISCUSSION

The emergence, rapid spread and international dissemination of the highly mutated Omicron variant raised concerns that this variant might soon become globally dominant and that several therapeutic or preventive interventions might be ineffective against this variant. The present study indicates that several of these concerns are justified. The S protein of the Omicron variant evaded antibody-mediated neutralization with higher efficiency than any previously analyzed S proteins of variants of interest and VOC. It was not appreciably inhibited by two antibody cocktails used for COVID-19 therapy and inhibition by antibodies induced by two immunizations with BNT was strongly reduced as compared to the spike of the Delta variant. Heterologous vaccinations and a BNT booster shot induced appreciable levels of neutralizing antibodies against the Omicron spike and might offer some protection against this variant. Nevertheless, our tools available to contain SARS-CoV-2 might require expansion and adaptation in order to efficiently combat the Omicron variant.

The findings that the Omicron spike facilitated efficient entry into several human cell lines and robustly bound human ACE2 suggest that the Omicron variant readily infects human cells. Nevertheless, some peculiarities of host cell entry driven by the Omicron spike relative to the B.1 spike were noted. The Omicron spike mediated slightly but significantly augmented entry into cell lines which only allow for cathepsin L- (293T, Huh-7, A549-ACE2) but not TMPRSS2-dependent entry, due to low or absent TMPRSS2 expression (Hoffmann *et al*., 2020b). We previously observed a similar phenotype for the S protein of another African SARS-CoV-2 lineage, A.30 (Arora et al., 2021a), and the mutations in spike responsible for this phenotype remain to be determined. Further, the Omicron spike used ACE2 proteins from different animal species with high efficiency for entry. The efficient usage of murine ACE2 is in keeping with the presence of mutations K417N and N501Y, which can also emerge upon SARS-CoV-2 adaptation to experimentally infected mice (Huang et al., 2021; Sun et al., 2021; Zhang et al., 2021b). In addition, the Omicron spike harbors mutations Q493R and Q498R that are related to exchanges Q493K and Q498H, which were also detected in mouse-adapted SARS-CoV-2 (Huang *et al*., 2021; Sun *et al*., 2021; Zhang *et al*., 2021b). However, some of these mutations are present in other VOC and cannot be taken as evidence that the Omicron variant may have evolved in infected mice in the wild. In addition, the Omicron spike was able to use ACE2 from the Pearson’s horseshoe bat (*Rhinolophus pearsonii*) with high efficiency. Again, this finding does not indicate the Omicron variant infects these animals (which are found in Asia) in the wild but rather reflects the ability of the Omicron spike to use diverse ACE2 orthologues for entry. Collectively, our results suggest that the mutations in the Omicron spike are compatible with robust usage of diverse ACE2 orthologues for entry and might thus have broadened the ability of the Omicron variant to infect animal species.

Cocktails of the antibodies Casirivimab and Imdevimab (REGN-COV2) as well as Etesevimab and Bamlanivimab (Eli Lilly) are used for COVID-19 therapy. Entry driven by the Omicron spike was fully or largely resistant against each of these antibodies and against the antibody cocktails, most likely due to mutations K417N, N440K, G446S, S477N, T478K, E484A, Q493R, G496S, Q498R, N501Y and Y505H, which are located within or close to the epitopes bound by these antibodies (Figure 2B). Thus, two frequently used COVID-19 treatment options will not be available to combat the Omicron variant. In contrast, Sotrovimab, a pan-sarbecovirus neutralizing antibody (Gupta *et al*., 2021), remained active against the Omicron spike, in keeping with Sotrovimab recognizing an epitope not substantially altered by mutations found in the Omicron spike (Figure 2B). Finally, soluble ACE2 robustly blocked entry driven by the Omicron spike and might be an option for treatment of patients infected with the Omicron variant.

Studies conducted before the emergence of the Omicron variant indicated that convalescent COVID-19 patients are efficiently protected against reinfection and antibody responses likely play an important role in protection (Addetia et al., 2020; Hall et al., 2021; Hansen et al., 2021; Harvey et al., 2021a; Krammer, 2021; Lumley et al., 2021). Our findings with serum/plasma samples collected in Germany during the first wave of the pandemic indicate that this high level of protection might not apply to reinfection with the Omicron variant. Thus, neutralization of the Omicron spike was 80-fold less efficient as compared to controls and several sera did not exert neutralizing activity. Although neutralization by sera from patients infected with the Delta variant remains to be examined, it is likely that convalescent patients might not be adequately protected against symptomatic reinfection with the Omicron variant, in keeping with recent data (Pulliam *et al*., 2021).

Neutralizing antibody responses are also believed to be critical for protection against COVID-19 by BNT vaccination (Corbett et al., 2021; Feng et al., 2021; Gilbert et al., 2021). We found that antibodies induced upon BNT/BNT immunization neutralized the Omicron spike with 34-fold reduced efficiency as compared to B.1. Therefore, two doses of BNT might not provide robust protection against severe disease induced by the Omicron variant and adaptation of the vaccine to the new variant seems required. In the meantime, heterologous AZ/BNT vaccination or a BNT booster (BNT/BNT/BNT) might afford some protection against the Omicron variant, since sera from individuals who had received the respective vaccinations neutralized the Omicron spike with appreciable efficiency. However, it remains to be determined whether this protection is transient or long lasting.

Our study has several limitations, including the use of pseudotyped virus and lack of analysis of T cell responses. However, given the important role that antibodies play in immune protection against SARS-CoV-2, our results suggest that preventive and therapeutic approaches have to be adapted for efficient protection against the Omicron variant. While such adaptations are in progress, heterologous or booster immunizations and conventional control measures like face masks and social distancing will help to limit the impact of the Omicron variant on public health.

## MATERIAL AND METHODS

### Cell culture

The following cells lines were used in the present study: 293T (human, female, kidney; ACC-635, DSMZ; RRID: CVCL_0063), A549-ACE2 ((Huang *et al*., 2021); based on parental A549 cells, human, male, lung; CRM-CCL-185, ATCC), BHK-21 (Syrian hamster, male, kidney; ATCC Cat# CCL-10; RRID: CVCL_1915, kindly provided by Georg Herrler, University of Veterinary Medicine, Hannover, Germany), Vero (African green monkey kidney, female, kidney; CRL-1586, ATCC; RRID: CVCL_0574, kindly provided by Andrea Maisner), Huh-7 cells (human, male, liver; JCRB Cat# JCRB0403; RRID: CVCL_0336, kindly provided by Thomas Pietschmann), Calu-3 (human, male, lung; HTB-55, ATCC; RRID: CVCL_0609, kindly provided by Stephan Ludwig) and Caco-2 cells (human, male, colon; HTB-37, ATCC, RRID: CVCL_0025). 293T, BHK-21, Vero and Huh-7 cells were maintained in Dulbecco’s modified Eagle medium (DMEM, PAN-Biotech). Calu-3 and Caco-2 cells were cultured in minimum essential medium (GIBCO) while A549-ACE2 cells were maintained in DMEM/F-12 medium (GIBCO). All media were supplemented with 10% fetal bovine serum (Biochrom) and 100 U/ml penicillin and 0.1 mg/ml streptomycin (PAA). Further, media for Calu-3 and Caco-2 cells were supplemented with 1x non-essential amino acid solution (from 100x stock, PAA) and 1 mM sodium pyruvate (GIBCO). Cell lines were validated by STR-typing, amplification and sequencing of a fragment of the cytochrome c oxidase gene, microscopic examination and/or according to their growth characteristics. In addition, all cell lines were regularly tested for mycoplasma contamination. For transfection of 293T cells, the calcium-phosphate based method was applied, while BHK-21 cells were transfected using Lipofectamine 2000 (Thermo Fisher Scientific).

### Plasmids

Plasmids encoding DsRed, VSV-G (vesicular stomatitis virus glycoprotein), soluble ACE2, SARS-CoV-2 S B.1 (codon optimized, contains C-terminal truncation of the last 18 amino acid) and SARS-CoV-2 S of VOCs Alpha (B.1.1.7), Beta (B.1.351), Gamma (P.1) and Delta (B. 1.617.2) have been previously reported (Brinkmann et al., 2017; Hoffmann et al., 2021a; Hoffmann et al., 2021b). The expression vector for the Omicron spike (based on isolate hCoV-19/Botswana/R40B58_BHP_3321001245/2021; GISAID Accession ID: EPI_ISL_6640919) was generated by Gibson assembly using five overlapping DNA strings (Thermo Fisher Scientific, sequences available upon request), linearized (BamHI/XbaI digest) pCG1 plasmid and GeneArt™ Gibson Assembly HiFi Master Mix (Thermo Fisher Scientific). Gibson assembly was performed according to manufacturer’s instructions. The pCG1 vector was kindly provided by Roberto Cattaneo (Mayo Clinic College of Medicine, Rochester, MN, USA). To generate of expression plasmids for different ACE2 orthologues (harboring a C-terminal cMYC-epitope tag) the respective sequences were amplified from existing plasmids (human, pangolin, cat, masked palm civet and red fox, *Rhinolophus landeri* ACE2) (Hoffmann *et al*., 2021b; Kruger et al., 2021; Li et al., 2021a) or ACE2 sequences were obtained from a commercial gene synthesis service (raccoon dog, American mink, Pig, Mouse, *Rhinolophus affinis, Rhinolophus sinicus, Rhinolophus pearsonii*; GeneArt). Expression plasmids for pangolin, masked palm civet and red fox ACE2 used as templates for cloning were kindly provided by Zhaohui Qian (Chinese Academy of Medical Sciences and Peking Union Medical College, Beijing, China). ACE2 open reading frames were inserted into the pQCXIP expression vector making use of NotI and PacI restriction sites. The integrity of all expression plasmids was confirmed using a commercial sequencing service (Microsynth SeqLab).

### Sequence analysis and protein models

S protein sequences and information on collection date and global distribution of SARS-CoV-2 isolates belonging to the Omicron variant were obtained from the GISAID (Global Initiative on Sharing All Influenza Data) database (https://www.gisaid.org). Structural models of the S protein were generated using YASARA software (http://www.yasara.org/index.html) and are based on a template that was constructed by modelling the SARS-2 S sequence on PDB: 6XR8 (Cai et al., 2020) using the SWISS-MODEL online tool (https://swissmodel.expasy.org). Alternatively, the following published crystal structures were employed; PDB: 6XDG (Hansen et al., 2020), PDB: 7L3N (Jones et al., 2021), PDB: 7C01 (Shi et al., 2020) or PDB: 6WPT (Pinto et al., 2020).

### Pseudotyping

Rhabdoviral pseudotypes harboring SARS-CoV-2 S protein were produced as described (Kleine-Weber et al., 2019). In brief, 293T cells were transfected with expression plasmids for SARS-CoV-2 S protein, VSV-G or empty plasmid (control). At 24 h posttransfection, cells were infected with at an MOI of 3 with a replication-deficient reporter VSV, termed VSV*ΔG-FLuc (Berger Rentsch and Zimmer, 2011). This virus lacks the ORF for VSV-G and codes for enhanced green fluorescent protein and firefly luciferase (FLuc). After 1 h of incubation, the input virus was removed and cells were washed with phosphate-buffered saline (PBS). Thereafter, culture medium containing anti-VSV-G antibody (culture supernatant from I1-hybridoma cells; ATCC no. CRL-2700) was added in order to neutralize residual inpunt virus. For cells expressing VSV-G, culture medium without antibody was added. After 16-18 h, the culture supernatant was harvested, separated from cellular debris by centrifugation for 10 min at 4,000 × g at room temperature (RT), and the clarified supernatants were aliquoted and stored at −80 °C.

### Analysis of spike protein-mediated cell entry

For analysis of S protein-driven cell entry, target cells were seeded in 96-well plates. In case of experiments addressing S protein usage of different ACE2 orthologues as receptor, cells were transfected with the respective ACE2 expression plasmids or empty vector. At 24 h post seeding (or transfection), the culture medium was removed and cells were inoculated with equal volumes of pseudotype preparations. At 16-18 h post inoculation, pseudotype entry was quantified by measuring the activity of virus-encoded luciferase in cell lysates. For this, cells were lysed using PBS supplemented with 0.5% triton X-100 (Carl Roth) for 30 min at RT. Subsequently, cell lysates were transferred into white 96-well plates, mixed with luciferase substrate (Beetle-Juice, PJK) and luminescence measured with a Hidex Sense Plate luminometer (Hidex).

### Analysis of ACE2 binding

The production of soluble ACE2 fused to the Fc portion of human immunoglobuline has been previously described in detail (Hoffmann *et al*., 2021b). In order to test binding of soluble ACE2 to the S protein, 293T cells seeded in 6-well plates were transfected with S protein expression plasmids or empty plasmid as negative control. At 24 h posttransfection, the medium was replaced with fresh culture medium. At 48 h posttransfection, the culture medium was aspirated and cells resuspended in PBS and pelleted by centrifugation at 600 × g for 5 min. Subsequently, cells were washed with PBS containing 1 % bovine serum albumin (BSA, PBS-B) and pelleted again. Next, the cell pellets were resuspended in 250 μl PBS-B containing soluble ACE2-Fc (1:100 dilution of 100x concentrated stock) and incubated for 60 min at 4 °C, employing a Rotospin test tube rotator disk (IKA). Thereafter, the cells were pelleted, resuspended in 250 μl PBS-B containing anti-human AlexaFlour-488-conjugated antibody (1:200; Thermo Fisher Scientific) and incubated again for 60 min at 4 °C. After a final wash step with PBS-B, the cells were fixed with 4 % paraformaldehyde solution for 30 min at RT, washed, resuspended in 150 μl PBS-B and analyzed by flow cytometry, using an LSR II flow cytometer and Flowing software version 2.5.1 (https://bioscience.fi/services/cell-imaging/flowing-software/).

### Analysis of inhibition of S protein-driven cell entry by soluble ACE2

S protein (or VSV-G) bearing particles were pre-incubated for 30 min at 37 °C with different dilutions of soluble ACE2 (undiluted, 1:10, 1:100, 1:1,000, 1:10,000). After incubation, the mixtures were added to Vero cells. Particles exposed to medium without soluble ACE2 served as control. Transduction efficiency was determined at 16-18 h postinoculation by determining luciferase activities in cell lysates, as described above.

### Collection of serum and plasma samples

Convalescent plasma/serum was obtained from COVID-19 patients treated at the University Medicine Göttingen (UMG) or Hannover Medical school (MHH).. Sera from individuals vaccinated with BNT162b2/BNT162b2 (BNT/BNT), ChAdOx1 nCoV-19/BNT162b2 (AZ/BNT) or BNT162b2/BNT162b2/BNT162b2 (BNT/BNT/BNT) were collected 14-72 days after receiving the last (second or third) dose at UMG or MHH (specific details on the samples can be found in Supplementary Table 1). All serum and plasma samples were heat-inactivated at 56 °C for 30 min. Collection of samples was approved by the ethic committee of the UMG (reference number: 8/9/20 and SeptImmun Study 25/4/19 Ü) and the Institutional Review Board of MHH (8973_BO_K_2020).

### Neutralization assay

For neutralization experiments, S protein bearing particles were pre-incubated for 30 min at 37 °C with Casirivimab, Imdevimab, Bamlanivimab, Etesevimab, Casirivimab + Imdevimab, Bamlanivimab + Etesevimab, Sotrovimab or unrelated control human IgG (5, 0.5, 0.05, 0.005, 0.0005 μg/ml). Alternatively, pseudotyped particles were pre-incubated with different dilutions of convalescent or vaccinated plasma/serum (dilution range: 1:25 to 1:12,800). After incubation, the mixtures were added to Vero cells. Particles exposed to medium without antibodies served as control. Transduction efficiency was determined at 16-18 h postinoculation by determining luciferase activities in cell lysates, as described above.

### Statistical analysis

The results on S protein-driven cell entry represent average (mean) data acquired from three to twelve biological replicates, each conducted with four technical replicates. The transduction was normalized against SARS-CoV-2 S B.1 (= 1). Alternatively, transduction was normalized against the background signal (luminescence measured for cells inoculated with particles bearing no viral glycoprotein; set as 1). For ACE2 binding, presented are the average (mean) geometric mean channel fluorescence data from six biological replicates, each conducted with single samples. The results on neutralization of spike protein-driven cell entry by monoclonal antibodies and IgG represent average (mean) data from three biological replicates (each conducted with technical quadruplicates) for which transduction was normalized against samples that did not contain any antibody (= 0% inhibition). The results on inhibition of spike protein-driven cell entry soluble ACE2 represent average (mean) data from three biological replicates (each conducted with technical quadruplicates) for which transduction was normalized against samples that did not contain soluble ACE2 (= 0% inhibition). The results on neutralization of spike protein-driven cell entry by convalescent plasma/serum or serum from vaccinated individuals are based on a single experiment, which was conducted with technical quadruplicates. For data normalization, the plasma/serum dilution factor that leads to 50% reduction in S protein-driven cell entry (neutralizing titer 50, NT50) was calculated. In addition, for each plasma/serum the fold change in NT50 between the indicated SARS-CoV-2 variants was calculated.

Error bars are defined as either standard deviation (SD) or standard error of the mean (SEM). Data were analyzed using Microsoft Excel (as part of the Microsoft Office software package, version 2019, Microsoft Corporation) and GraphPad Prism 8 version 8.4.3 (GraphPad Software). Statistical significance was analyzed by two-tailed Student’s t-test with Welch correction (pseudotype entry, ACE2 binding), two-way analysis of variance (ANOVA) with Dunnett’s or Sidak’s post-hoc test (soluble ACE2 inhibition, and monoclonal antibody neutralization) or Wilcoxon matched-pairs signed rank test (serum/plasma neutralization). Only p-values of 0.05 or lower were considered statistically significant (p > 0.05, not significant [ns]; p ≤ 0.05, *; p ≤ 0.01, **; p ≤ 0.001, ***). Details on the statistical test and the error bars can be found in the figure legends.

## SUPPLEMENTAL INFORMATION

**Figure S1:** Cell entry driven by the spike proteins of VOC (related to Figure 1F).

Pseudotype entry data presented in Figure 1F normalized against the assay background. The experimental set-up is identical as described in the legend of Figure 1 (panel F) with the difference that pseudotype entry was normalized against signals obtained from cells inoculated with particles bearing no viral glycoprotein (background, set as 1) and inclusion of data produced for particles bearing VSV-G. Error bars indicate the standard error of the mean.

**Figure S2:** Individual neutralization data (Related to Figure 3).

Presented are the individual neutralization results for the data shown in Figure 3. Data represent the mean values of four technical replicates with error bars indicating the standard deviation. The curves were calculated based on a non-linear regression model with variable slope.

## ACKNOWLEDGMENTS

S.P. acknowledges funding by BMBF (01KI2006D, 01KI20328A, 01KI20396, 01KX2021), the Ministry for Science and Culture of Lower Saxony (14-76103-184, MWK HZI COVID-19) and the German Research Foundation (DFG; PO 716/11-1, PO 716/14-1). M.S.W. received unrestricted funding from Sartorius AG, Lung research. H.-M.J. received funding from BMBF (01KI2043, NaFoUniMedCovid19-COVIM: 01KX2021), Bavarian State Ministry for Science and the Arts and Deutsche Forschungsgemeinschaft (DFG) through the research training groups RTG1660 and TRR130. G.B. acknowledges funding by German Center for Infection Research (grant no 80018019238).

## AUTHOR CONTRIBUTIONS

Conceptualization, M.H., S.P.; Funding acquisition, H.M.J., G.B., S.P.; Investigation, M.H., N.K., C.R., A.K., I.N., L.G., A.-S.M., Essential resources, S.S., A.C., M.S.W., M.L., A.D.-J., H.-M.J., G.M.N.B.; Writing, M.H., S.P., Review and editing, all authors.

