## Supplementary figures and images for "The Omicron variant is highly resistant against antibody-mediated neutralization – implications for control of the COVID-19 pandemic"

### Supplemental figure S1

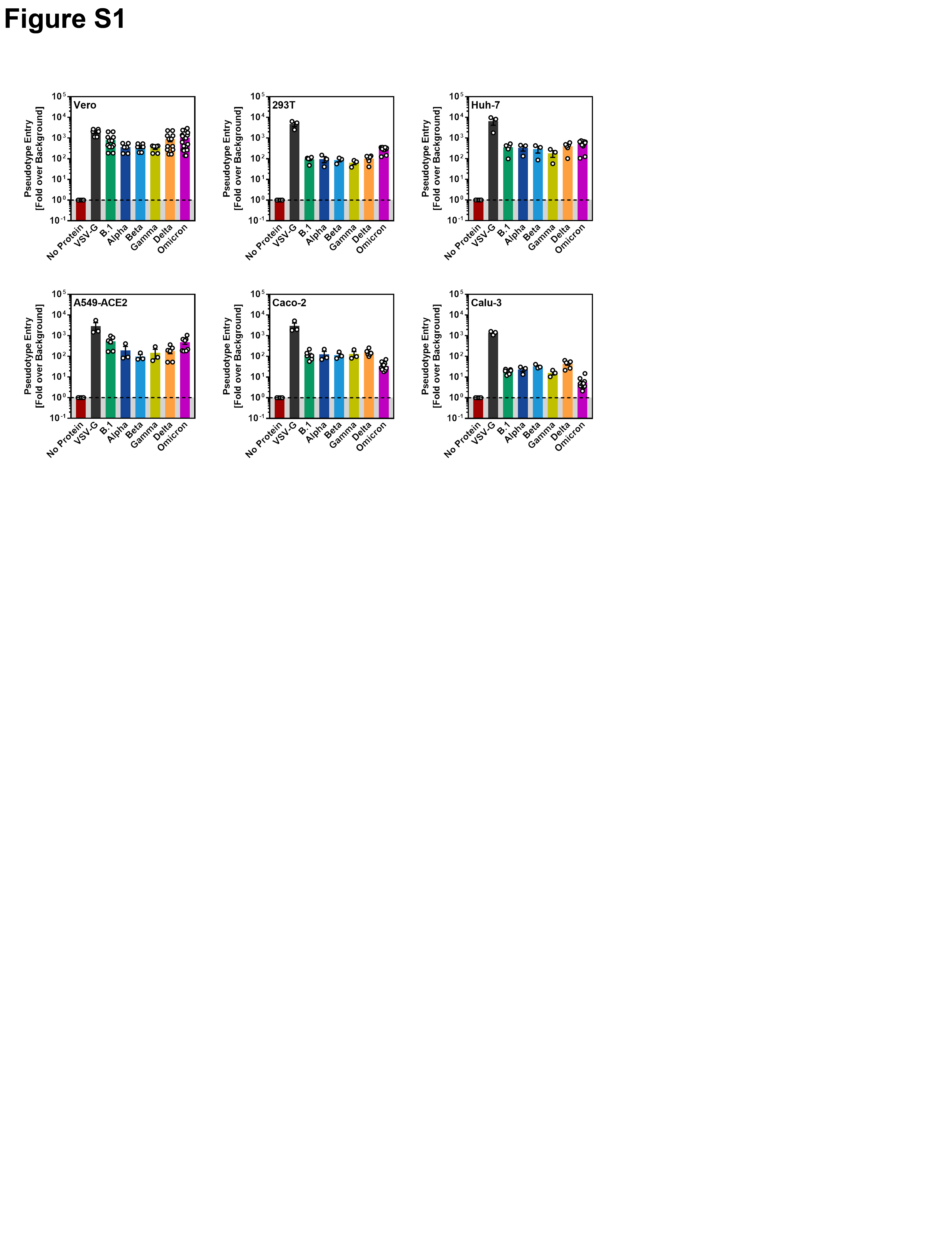

### Supplemental figure S2

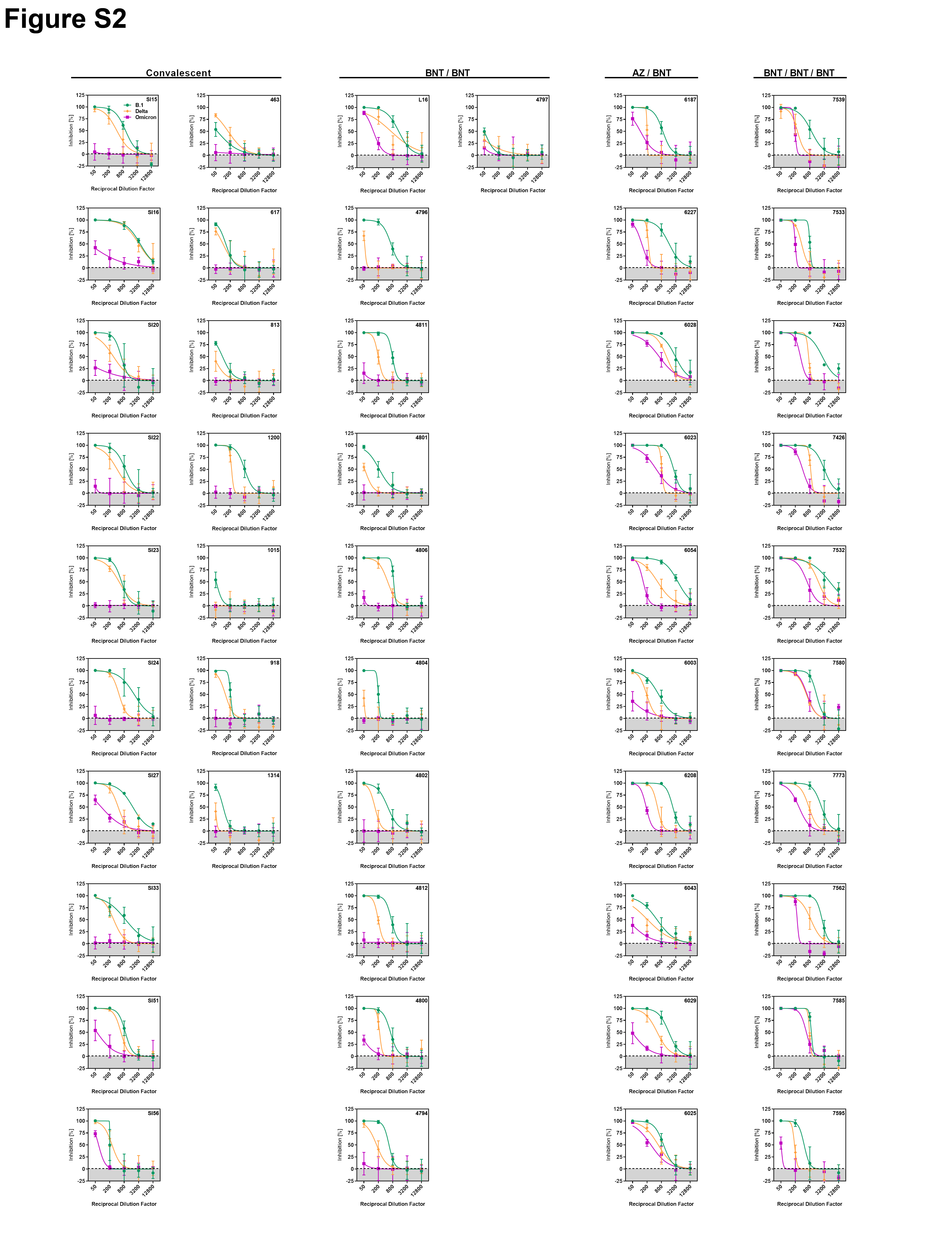
