## Supplemental table S1 for "The Omicron variant is highly resistant against antibody-mediated neutralization – implications for control of the COVID-19 pandemic"

**COVID-19 patient data (related to Figure 3)**

| **Identifier** | **Type** | **Period** | **Symptoms** |
| --- | --- | --- | --- |
| **SI15** | Convalescent | 1st wave | Severe* |
| **SI16** | Convalescent | 1st wave | Severe* |
| **SI20** | Convalescent | 1st wave | Severe* |
| **SI22** | Convalescent | 1st wave | Severe* |
| **SI23** | Convalescent | 1st wave | Severe* |
| **SI24** | Convalescent | 1st wave | Severe* |
| **SI27** | Convalescent | 1st wave | Severe* |
| **SI33** | Convalescent | 1st wave | Severe* |
| **SI51** | Convalescent | 1st wave | Severe* |
| **SI56** | Convalescent | 1st wave | Severe* |
| **463** | Convalescent | 1st wave | Mild |
| **617** | Convalescent | 1st wave | Mild |
| **813** | Convalescent | 1st wave | Mild |
| **1200** | Convalescent | 1st wave | Mild |
| **1015** | Convalescent | 1st wave | Severe* |
| **918** | Convalescent | 1st wave | Mild |
| **1314** | Convalescent | 1st wave | Mild |

*: admission to intensive care unit required

**Table S2**

**Vaccinated patient data (related to Figure 3)**

| **Identifier** | **Type** | **Vaccination** | **Age group (years)** | **Documented SARS-CoV-2 infection?** | **Time since last vaccination (days)** |
| --- | --- | --- | --- | --- | --- |
| **4796** | Vaccinated | BNT/BNT | 35-44 | none | 14 |
| **4811** | Vaccinated | BNT/BNT | 25-34 | none | 15 |
| **4801** | Vaccinated | BNT/BNT | 25-34 | none | 18 |
| **4806** | Vaccinated | BNT/BNT | 25-34 | none | 19 |
| **4804** | Vaccinated | BNT/BNT | 55-64 | none | 21 |
| **4802** | Vaccinated | BNT/BNT | 45-54 | none | 21 |
| **4812** | Vaccinated | BNT/BNT | 35-44 | none | 21 |
| **4800** | Vaccinated | BNT/BNT | 45-54 | none | 21 |
| **4799** | Vaccinated | BNT/BNT | 25-34 | none | 21 |
| **4797** | Vaccinated | BNT/BNT | 35-44 | none | 22 |
| **L16** | Vaccinated | BNT/BNT | 18-24 | none | 72 |
| **6187** | Vaccinated | AZ/BNT | 25-34 | none | 14 |
| **6227** | Vaccinated | AZ/BNT | 25-34 | none | 14 |
| **6208** | Vaccinated | AZ/BNT | 55-64 | none | 14 |
| **6043** | Vaccinated | AZ/BNT | 45-54 | none | 14 |
| **6029** | Vaccinated | AZ/BNT | 35-44 | none | 14 |
| **6025** | Vaccinated | AZ/BNT | 35-44 | none | 14 |
| **6028** | Vaccinated | AZ/BNT | 35-44 | none | 14 |
| **6023** | Vaccinated | AZ/BNT | 55-64 | none | 14 |
| **6054** | Vaccinated | AZ/BNT | 45-54 | none | 14 |
| **6003** | Vaccinated | AZ/BNT | 35-44 | none | 14 |
| **7539** | Vaccinated | BNT/BNT/BNT | 25-34 | none | 14 |
| **7533** | Vaccinated | BNT/BNT/BNT | 35-44 | none | 17 |
| **7423** | Vaccinated | BNT/BNT/BNT | 25-34 | none | 19 |
| **7426** | Vaccinated | BNT/BNT/BNT | 55-64 | none | 19 |
| **7532** | Vaccinated | BNT/BNT/BNT | 45-54 | none | 19 |
| **7580** | Vaccinated | BNT/BNT/BNT | 45-54 | none | 20 |
| **7773** | Vaccinated | BNT/BNT/BNT | 35-44 | none | 21 |
| **7562** | Vaccinated | BNT/BNT/BNT | 25-34 | none | 24 |
| **7585** | Vaccinated | BNT/BNT/BNT | 45-54 | none | 24 |
| **7595** | Vaccinated | BNT/BNT/BNT | 55-64 | none | 26 |
